# Epigenetic-structural changes in X chromosomes promote Xic pairing during early differentiation from mouse embryonic stem cells

**DOI:** 10.1101/2021.10.25.465811

**Authors:** Tetsushi Komoto, Masashi Fujii, Akinori Awazu

## Abstract

X chromosome inactivation center (Xic) pairing is robustly observed during the differentiation of embryonic stem (ES) cells from female mouse embryos, and this process is related to X chromosome inactivation, the circadian clock, intra-nucleus architecture, and metabolism. However, the mechanisms underlying the identification and approach of X chromosome pairs in the crowded nucleus are unclear. To elucidate the driving force of Xic pairing, we developed a coarse-grained molecular dynamics model of intranuclear chromosomes in ES cells and in cells 2 days after the onset of differentiation (2-days cells) by considering intrachromosome epigenetic-structural feature-dependent mechanics. The analysis of the experimental data showed X-chromosomes change to specifically softer than autosomes during the cell differentiation by the rearrangement of their distributions of open-close chromatin regions, and the simulations of these models exhibited such softening promoted the mutual approach of the Xic pair. These findings suggested that local intrachromosomal epigenetic features may contribute to the regulation of cell species-dependent differences in intranuclear architecture.

## Introduction

The pairing between X chromosomes, particularly that between the regions called X chromosome inactivation center (Xic) regions, is robustly observed during the early differentiation of embryonic stem (ES) cells from female mouse embryos [1–4]. Xic pairing is expected to play important roles in X chromosome inactivation (XCI) and may link mechanisms regulating intranuclear architecture, metabolic programs, and the onset of the circadian clock [4]. Additionally, understanding of the physiological and biophysical mechanisms of XCI-related X chromosome behaviors in mouse embryos may provide insights into the mechanisms of XCI in various mammals, including humans [5]. Indeed, various models have recently been proposed for postpairing processes of X chromosomes, such as XCI promotion and Xic pairing stabilization [6–12], which are expected to occur in the inter-X chromosome compartment with a width ∼100 *nm* [13]. However, the mechanisms through which X chromosome pairs can find and approach each other in a large, crowded nucleus (diameter ∼10 μm, containing 40 total chromosomes) have not yet been clarified.

In studies using mouse ES cells, various epigenomic features, such as histone modification distributions and positions of chromatin domain boundaries in the X chromosome, show major differences among cells at the onset of differentiation and at 2 days after; subsequently, X chromosome pairing can be observed [2–4]. Recent studies of intranuclear chromosome structural dynamics have suggested that genomic features such as chromosome size and the ratio of transcribed genome regions, and epigenomic features such as histone modifications and specific protein binding, may influence intranuclear chromosomes and intrachromosome chromatin domain positioning [14–28]. Additionally, some theoretical studies based on polymer chains with heterogeneities assuming genomic and epigenomic feature-dependent physical properties of local chromatin succeeded in reproducing experimentally observed nucleus-wide genome structures, such as transcriptionally active/inactive chromatin distributions in interphase of human cells [24] and mouse rod cell [25, 27], as well as the pairing of homologous chromosomes in meiotic prophase of fission yeast [28]. These facts suggest that epigenomic state changes in X chromosomes may promote Xic pairing during the differentiation of mouse ES cells.

Accordingly, in this study, we developed coarse-grained molecular dynamics models of nuclei (containing 40 chromosomes) from mouse ES cells (the ES cell model) and cells at 2 days after the start of differentiation (the 2-days cell model) to test this hypothesis. Chromosomes were described by blob chains with heterogeneity of physical properties based on their respective high-throughput chromosome conformation capture (Hi-C) data [29].

## Results

### A/B-compartment distributions of X chromosomes in ES cells were unstable but stabilized during differentiation

To evaluate chromosome states, A/B-compartments of ES cells and cells at 2 days after the onset of differentiation (2-days cells) were evaluated using Hi-C data (GSM3127755, GSM3127759, GSM3127756, GSM3127760) and GC-content of local chromatin regions of mouse genome (mm9) [29]. Here, regions corresponding to open chromatin were labeled as A-compartment regions, whereas those corresponding to closed chromatin were labeled as B-compartment regions [30]. In ES cells, the profile of A/B-compartment locations in X chromosomes showed large deviations among the results from two biological replications, although those of autosomes were robust (Fig 1a). Accordingly, the genomic region in which both replicates exhibited A-compartments or B-compartments was defined as the A-region or B-region, respectively, whereas the genomic region in which different compartments were obtained among the results from two replicates was defined as the M-region.

**Fig 1.**
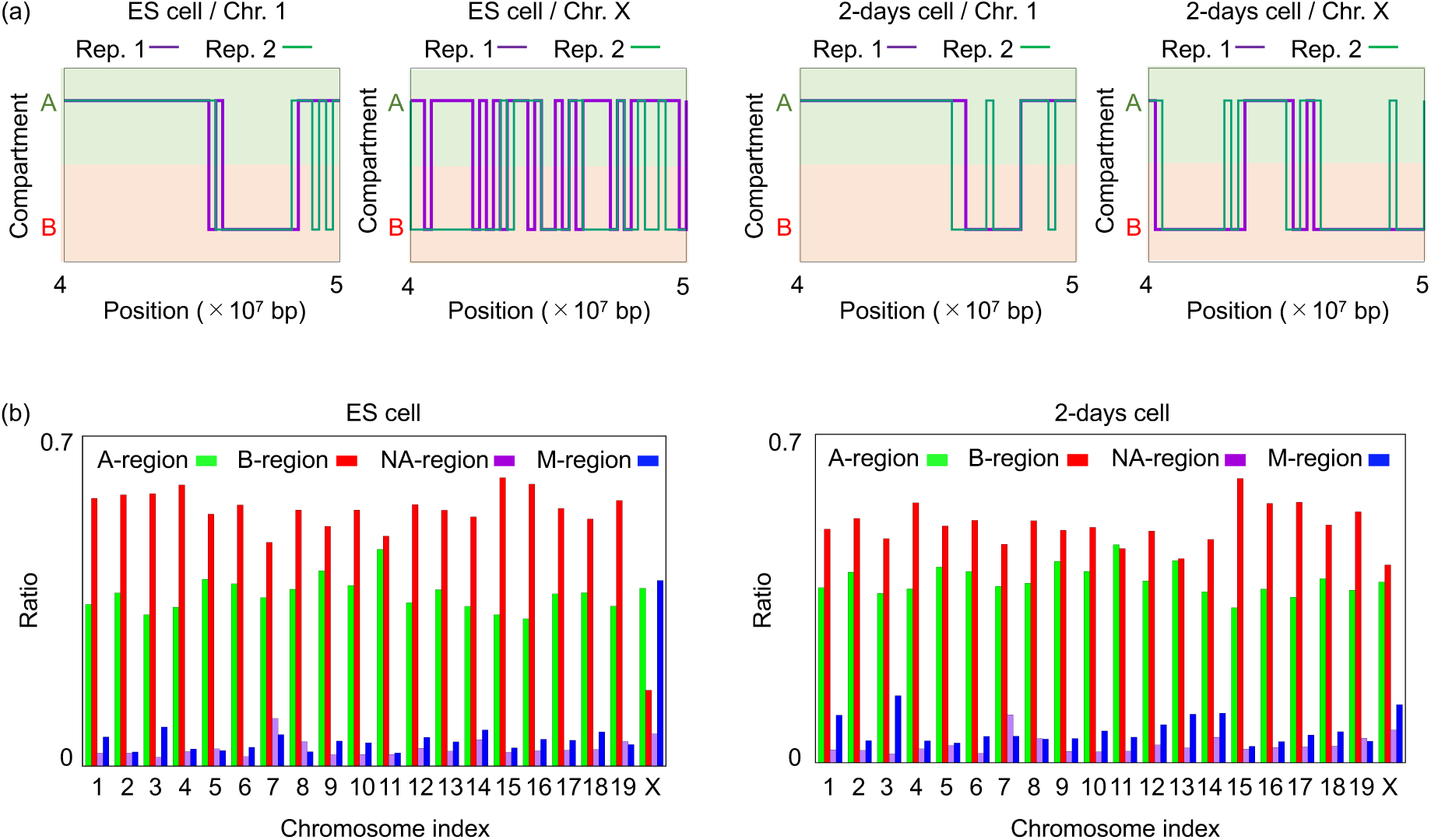
Intragenomic distributions and ratios of A-, B-, and M-regions on autosomal and X chromosomes in ES cells and 2-days cells: (a) Example distribution profiles of A/B compartments in two biological replicates and those of A-, B-, and M-regions on chromosomes 1 and X (4 × 10^7^ bp to 5 × 10^7^ bp). The M-regions were broadly distributed on X chromosomes of ES cells but showed decreased distributions in 2-days cells. (b) Ratios of A-, B-, and M-regions on each chromosome. Both ES cells and 2-day differentiated cells contained few M-regions on autosomes. The ratio of the M-region on the X chromosome was larger in ES cells and dramatically reduced in 2-days cells. The NA region is the region with no Hi-C data, which was expected to correspond to the region containing the centromere or telomere.

The M-region was distributed broadly and occupied the largest genomic region among X chromosomes from ES cells (Fig 1b). The epigenetic state of the M-region was expected to change temporally and to exhibit intermediate features of the A- and B-regions on average. By contrast, in 2-days cells, the M-region was dramatically reduced across the entire chromosome, whereas the B-region was increased (Fig 1b). Compared with the X chromosome, the intrachromosome occupation of the A-, B-, and M-regions did not exhibit large variations in autosomes during this cell differentiation process.

### Spatial distributions of open/closed chromatin regions on the X chromosome changed following differentiation

Next, the spatial distributions of A-, B-, and M-regions in each chromosome in ES cells and 2-days cells were measured based on previous results obtained from Hi-C data [29] (https://doi.org/10.5281/zenodo.3371884) and the following procedure. First, the basic conformation of each chromosome was evaluated as a polymer by analysis of the Hi-C contact matrices. Here, the position of each monomer in the obtained polymer provided the intrachromosome spatial position of each region (Table S1). Second, the number distributions of monomers with A-, B-, and M-regions as a function of the distance from the center of each chromosome to these monomers (*DCC*) were evaluated and were, respectively, designated *ND*_*A*_, *ND*_*B*_, and *ND*_*M*_ (Fig 2a). *ND*_*α*_ was defined as the number of monomers with *α* -region between *DCC* − 0.05 (μm) and *DCC* + 0.05 (μm). Third, the radial distributions of A-, B-, and M-regions, designated *RD*_*A*_, *RD*_*B*_, and *RD*_*M*_, respectively, and the radial probability distributions of monomers, *RPD*, for each chromosome were evaluated as a function of *DCC*, where *RD*_*X*_ and *RPD* were defined as *RD*_*X*_ = *ND*_*X*_/4*πDCC*^2^ (*X* = *A, B, or M*) and *RPD* = (*ND*_*A*_ + *ND*_*B*_ + *ND*_*M*_)/4*πDCC*^2^*N*_*C*_ (*N*_*C*_ = the number of monomers in each chromosome), respectively.

**Fig 2.**
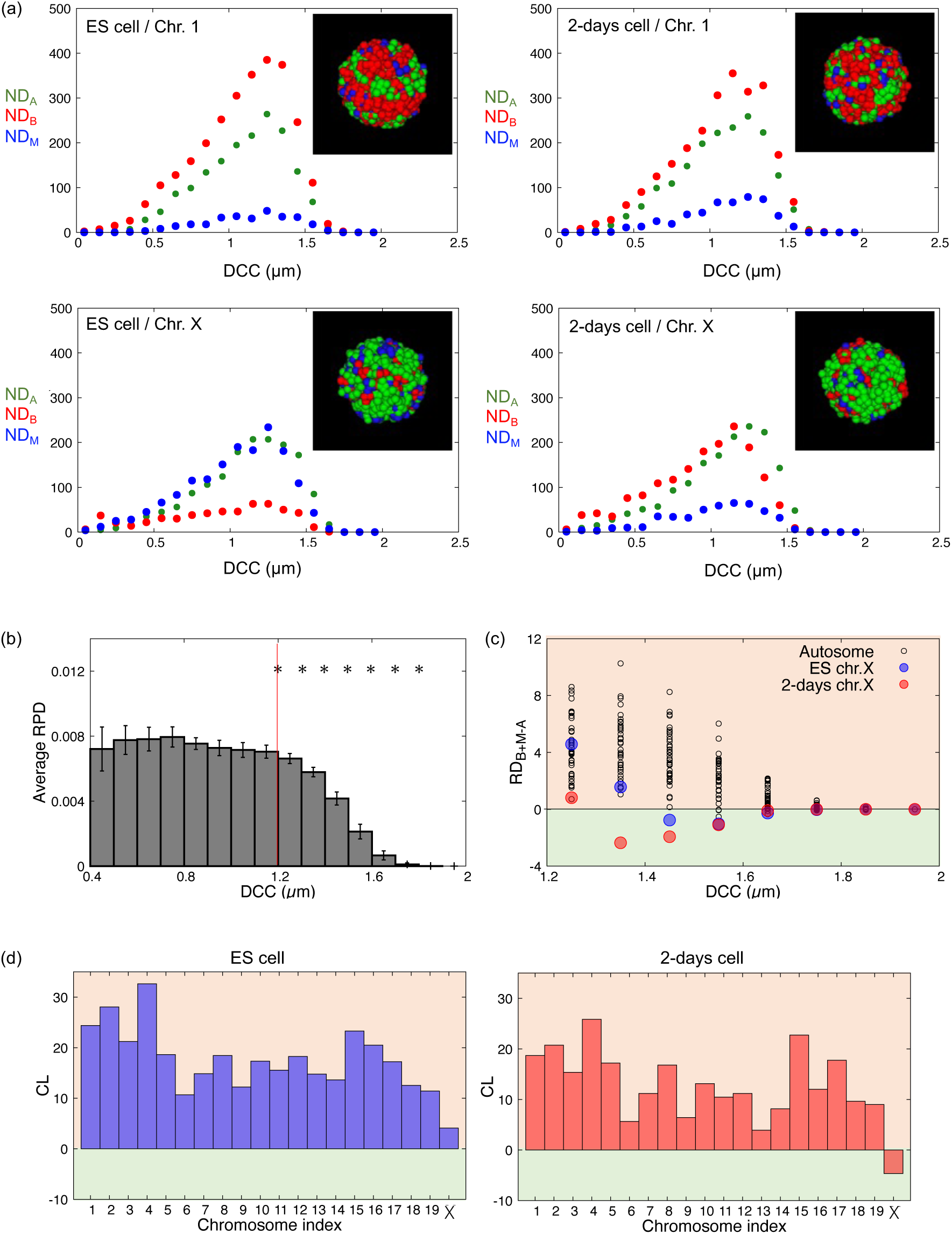
Intrachromosome spatial distributions of open and closed chromatin regions. (a) *ND*_*A*_, *ND*_*B*_, and *ND*_*M*_, of chromosome 1 and the X chromosome in ES cells and 2-days cells. Insets show typical distributions of A-(green), B- (red), and M-regions (blue) on chromosome surfaces, as observed from one viewpoint (see data in Table S1). (b) Average *RPD* values and 95% confidence intervals (error bars) for all chromosomes. Stars between two neighboring *DCC* values indicated that two average *RPD* values were significantly different (*p* < 0.01 using Welch’s test; *p* values are plotted in Fig S1). (c) *RD*_*B*+*M* − *A*_ at *DCC* > 1.2 for each chromosome. (d) *CL* for each chromosome.

The *RPD*s of chromosomes in both ES cells and 2-days cells exhibited dramatic decreases, with *DCC* values greater than ∼1.2, where the average *RPD* over all chromosomes showed a significant decrease (*DCC* ≥ 1.25; Fig 2b, Fig S1). Accordingly, monomers showing values of *DCC* ≥ 1.25 were expected to be localized on the surface of each chromosome.

For *DCC* ≥ 1.25, *RD*_*B* +*M* − *A*_ = *RD*_*B*_ + *RD*_*M*_ − *RD*_*A*_ was measured as a function of *DCC* (Fig 2c), and the chromatin closedness of each chromosome surface (*CL* = ∑_*DCC >*1.2_ *RD*_*B* +*M* − *A*_) was evaluated in both ES cells and 2-days cells (Fig 2d). From the results, we observed that only X chromosomes in 2-days cells exhibited negative *CL* values, indicating that the surface of the chromosome was predominantly occupied by open chromatin regions, although other chromosomes were predominantly surrounded by closed or potentially closed chromatin regions.

### Simulations of the coarse-grained blob chain model showed mutual approach of Xic pairs in differentiated cells

Simulations of models of intranuclear chromosome dynamics in ES cells (ES cell model) and 2-days cells (2-days cell model) were performed using coarse-grained blob chain models. These models were constructed using the previously obtained polymers that gave the basic conformations of chromosomes and the distributions of A-, B-, and M-regions. The populations of monomers successively connected along the polymer were divided into blobs containing local chromatin parts with around 0.1–10 Mbp based on estimations of boundary scores [29] and appropriate chromosome-dependent characteristic distances along genome sequences. The coordinate and radius of each blob was defined as the centroid coordinate of the blob containing monomers and the standard deviation of the distance from the blob centroid to the monomers, where blob radii were approximately 0.15–1 μm. Such chains of blobs provided the basic structure for the coarse-grained model of chromosomes, showing an envelope shape similar to that of the original polymer by elimination of small structural noises expected to be due to the noise of Hi-C data. Additionally, the epigenetic attributes of A, B, or M were added to each blob based on our other findings (Fig 3a, b).

**Fig 3.**
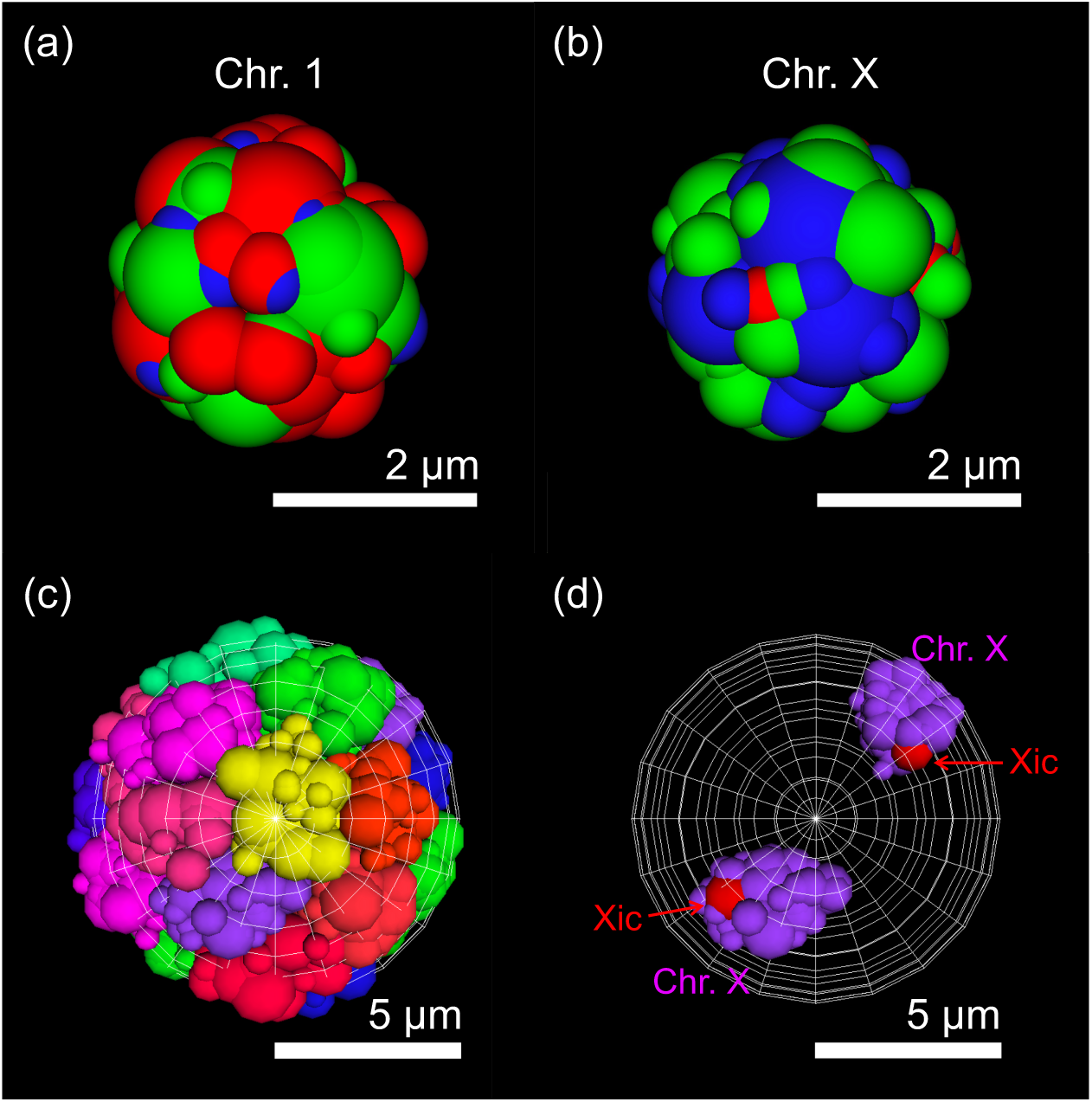
Snapshots of simulations of the coarse-grained blob chain model of chromosomes. (a, b) Examples of snapshots of blob chain models of chromosome 1 (a) and the X chromosome (b) in the ES cell model (see Table S2 for results for other chromosomes in both the ES cell and 2-days cell models). Green, red, and blue spheres indicate A-blob, B-blob, and M-blob, respectively. (c, d) Snapshot of all chromosomes in the ES cell model (c) and positioning of two X chromosomes (d). Blob chains with different colors indicate models of different chromosomes.

Each simulation was performed by considering the force required to maintain the basic structure of blob chains and the repulsion between blobs due to their excluded volumes. The A-blob consisted predominantly of open chromatin regions, which were sparse and flexible, whereas the B-blob consisted predominantly of closed chromatin regions, which contained condensed nucleosomes and proteins. Thus, the rigidity of the blob was assumed to increase in the order A < M < B. The nuclear membrane was assumed to be a spherical shell with a radius of 5 μm (Fig 3c, d).

Thirty simulations were performed for both the ES cell model and the 2-days cell model from different initial configurations, and the time course and probability distributions of the distance between two blobs containing Xic regions were plotted (Fig S2, Fig 4a). Significantly higher probability values at small distances and significantly smaller probability values at large distances were obtained in the 2-days cell model compared with those in the ES cell model. This result indicated that two blobs with Xic regions in the 2-days cell model tended to approach each other significantly more frequently than those in the ES cell model, consistent with experimental observations [1–4]. The probability distributions of the distance between the pair of X chromosomes also exhibited similar model-dependent features (Fig 4b). By contrast, for most homologous autosomes, the features of the probability distribution of the distances did not exhibit significant changes between ES cell and 2-days cell models (Fig 4c, Fig S3).

**Fig 4.**
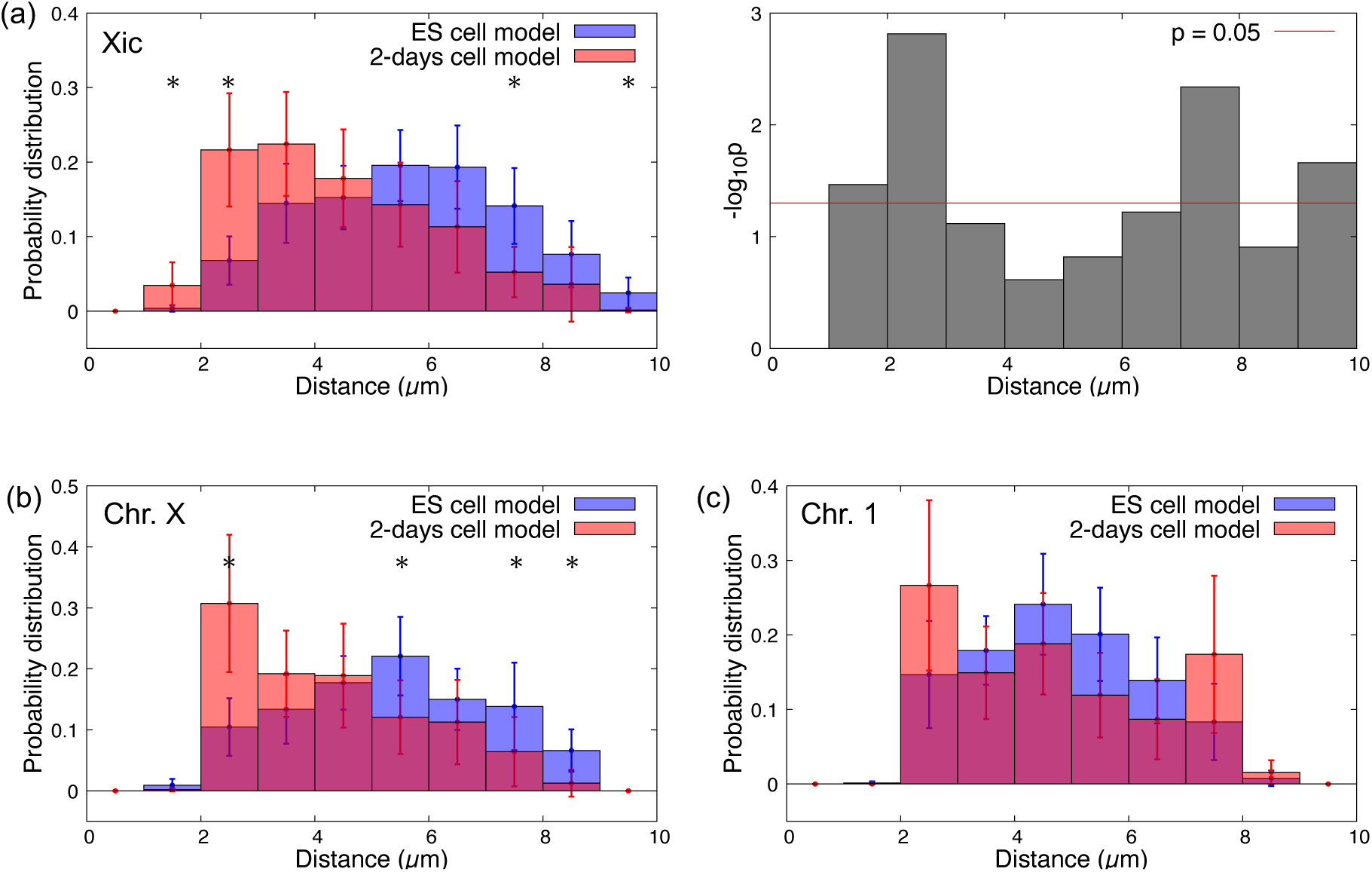
Probability distributions of the distance between the Xic pair and between homologous chromosomes. (a) Average and 95% confidence interval (error bar) of probability distributions of the distance between the pair of blobs containing Xic in the ES cell and 2-days cell models (left), and *p* values (Welch’s test) between two probability values of the two models at each distance (right). (b, c) The same plots as the left for (a) between an X chromosome pair (b) and between a chromosome 1 pair (c). The average and 95% confidence interval were evaluated using results from 30 simulations for each model. Distances with asterisks indicate that differences in probability values between the two models were significant (*p* < 0.05 using Welch’s test; see also Figs S3 and S4).

## Discussion

From simulations of the coarse-grained model of intranuclear chromosome dynamics, we found that two X chromosomes approached each other more frequently in the 2-days cell model than in the ES cell model. This result was consistent with recently reported experiments [1–4].

Now, we will consider the driving force that promotes this intranuclear phenomenon based on the mechanical properties of chromosomes given by their intrastructural epigenetic features. Analysis of Hi-C data revealed that the X chromosome showed dramatic changes in its epigenetic state during the differentiation process, as follows. The chromatin region, for which the epigenetic state was not determined, was stably distributed broadly in X chromosomes in ES cells. These unstable regions could also be regarded as potential closed chromatin regions. By contrast, in all chromosomes in 2-days cells, most chromatin regions were stably open or closed. Additionally, both open chromatin regions and (true and potential) closed chromatin regions were also located at the surface of the X chromosome in ES cells, whereas only an open chromatin dominant region was located at the surface of the X chromosome in 2-days cells. Autosomes exhibited minor changes in such structural epigenetic states during differentiation.

In 2-days cells, X chromosomes were predominantly surrounded by open chromatin regions, whereas the surfaces of autosomes were predominantly occupied by closed chromatin regions. Thus, X chromosomes and autosomes may be regarded as soft and rigid objects, respectively, and repulsion caused by the excluded volume between X chromosomes was much weaker than that between other chromosome pairs. Therefore, the pair of adjoining X chromosomes could form a more compact shape, providing a larger intranuclear space for the other chromosomes. Thus, the number of states correlating with the entropy of the system was expected to be larger when two X chromosomes were spatially close than when other chromosomes were located between the X chromosomes. This entropic effect was expected to be the driving force for the mutual approach of X chromosomes, similar to the force known as depletion force [31–33] or to that inducing the phase separation of polymers with different rigidities [22, 34–36].

In ES cells, the large area of the X chromosome surface was occupied by closed and potentially closed chromatin regions. Thus, the excluded volume effect between X chromosomes in ES cells was not higher than that between other chromosome pairs but was sufficiently higher than that between X chromosomes in 2-days cells. Based on this, we concluded that the mutual approach of X chromosomes occurred less frequently in ES cells than in 2-days cells because of weakening of this driving force owing to entropic effects.

The main purpose of this study was exploration of the driving force of nucleus-wide chromosome migration, resulting in X chromosome pairing, which preceded the local shortrange interaction between two Xic pairs through specific molecules, such as Xist and Tsix RNAs. Thus, any effects of Xist RNA were not considered explicitly. From a combination of the current model and other recently proposed models of Xic interactions based on microscopic molecular processes, however, a model may be developed to explain all processes involved in Xic pairing and stabilization.

Additionally, recent experimental data have demonstrated that Xic tends to localize to the nuclear periphery owing to the association between lamin-binding receptors (Lbrs) and Xist RNA; this tendency is weakened by inhibition of Xist RNA [37]. Because our current model did not consider the effects of such RNA, Xic did not exhibit strong specific localization to the nuclear periphery (Fig S5), consistent with recent experiments. In future studies, a modified model with nontrivial interactions between Xic and the nuclear periphery through the Lbr-Xist association and the Xist-Tsix-driven inter-Xic interaction should be developed to consider the entirety of Xic behaviors during the early differentiation process of ES cells.

Overall, our current findings suggested that intrachromosomal structural epigenetic features and their changes played important roles in regulating intranuclear positioning and transitions. Application of such arguments is expected to provide important insights into the mechanisms underlying cell species- and cell cycle-dependent behaviors of intranuclear architecture.

## Model and Methods

### Determination of A-, B-, and M-regions in chromosomes from ES cells and 2-days cells

A- and B-compartments for the first and second replicates of ES cells and 2-days cells were obtained as previous proposed, using Hi-C data (GSM3127755, GSM3127759, GSM3127756, GSM3127760) and GC-content of local chromatin regions of mouse genome (mm9) [29].

Each genome region was defined as an A- or B-region if the region was labeled as an A- or B-compartment in the results from two biological replicates of each experiment. The genome region was defined as an M-region if the region was labeled as a different compartment in results from two replicates for each experiment. Regions with no Hi-C data were defined as NA regions and were expected to correspond to centromere or telomere regions.

### Estimation of the basic structures of chromosomes using polymer models from Hi-C data and radial distributions of each region

A polymer model of the average three-dimensional structure of each chromosome was constructed using Hi-C data for ES cells and 2-days cells (https://doi.org/10.5281/zenodo.3371884) [29] based on a recently proposed consensus method [38–40] implemented in PASTIS as an MDS2 method [41]. For both ES and 2-days cells, merged Hi-C data obtained from two biological replicates were analyzed.

The three-dimensional spatial position of each monomer in each polymer model for each chromosome provided that of the corresponding genome region analyzed in the previous section (Table S1). Based on our results, each monomer was attributed to the A-, B-, or M-region (Fig S6a).

According to these position data, *ND*_*A*_, *ND*_*B*_, and *ND*_*M*_ were evaluated in ES and 2-days cells (Fig 2). The results were confirmed to be unchanged qualitatively, even if the three-dimensional structure of each chromosome was constructed using other methods implemented in PASTIS with NMDS, PM1, and PM2 methods [41] (data not shown).

### Determination of A-, B-, and M-domains and domain groups in each chromosome

Each polymer model for each chromosome was divided into groups of successively connected monomers, called domains (Fig S6a). Monomers acting as domain boundaries (boundary monomers) were obtained based on boundary scores that could be calculated from merged Hi-C data using a previously proposed method with the same parameter values (Table S1) [29]. In this study, each domain was assumed to consist of one boundary monomer and the monomers between this boundary monomer and the upstream boundary monomers (Fig S6a).

The epigenetic attribute of each domain was defined as the A-domain if more than 50% of the domain occupied the A-region, as the B-domain if more than 50% of the domain occupied the A-region, or as the M-domain if the domain was neither an A-domain nor a B-domain (Fig S6a). Notably, the NA region was considered a B-region when determining the domain attributes because the NA region was expected to correspond to the centromere or telomere regions forming heterochromatin. Additionally, the domain group was defined as the group containing domains with the same epigenetic attributes successively connected along the polymer (Fig S6a).

### Description of each domain group using blobs to construct coarse-grained models of chromosomes

Each domain group was divided into blobs, forming a similar envelope shape as the monomer population in the blob (Fig S6b), with elimination of small structural noise as follows (Fig S6a).

First, k-means clustering of the three-dimensional coordinates of monomers in each domain group was performed. For this clustering, the number of clusters dividing each domain group in the n-th chromosome was assumed as the integer value with [number of domains in the domain group] / CND_n_ truncated, where CND_n_ was the characteristic number of domains in the n-th chromosome and was defined as the number of domains for which the physical distance between two loci increased successively on average with the increases in the domain number between the two loci. By this definition, each domain group containing more domains than CND_n_ was divided into multiple clusters, enabling imitation of its curved shape by connection of these cluster centers.

CND_n_ was evaluated by comparing the decay rate of the average PDND ([physical distance between two domains] / [number of domain connections between these domains]) as a function of the number of domain connections between the two domains. Because the average PDND in each chromosome decreased approximately as ∼ exp(-[number of domains between two loci] / L) (Fig S7), CND_n_ was assumed to be L.

Next, each domain group was described by distinct blobs based on the results of clustering. If the monomer cluster consisted of only a set of monomers successively connected along the polymer, the set of monomers was described as one blob. If not, the monomer cluster was divided to subclusters in which monomers were successively connected along the polymer in each subcluster, and each set of monomers in each subcluster was described as one blob.

The coordinate and radius of each blob were defined as the centroid coordinate of blobs containing monomers and the standard deviation of the distance from the blob centroid to the contained monomers. The epigenetic attribute of each blob was given by that of the corresponding domain group and designated the A-, B-, and M-blobs. By connecting the neighboring blobs along the polymer, a coarse-grained model of chromosomes was constructed as blob chains involving structural, genomic, and epigenomic features (Fig S6a, Table S2). In this model, the distance between each pair of blobs in the same chain was defined as that in the basic structure of the blob chain model of chromosomes.

### Equation of motion for each blob in the coarse-grained model of chromosomes

The dynamics of all intranuclear chromosomes (chromosomes 1–19 and the X chromosomes) of ES cells and 2-days cells were simulated using the motion of each blob (Table S2), which was affected by the interaction potential and noise from the nucleoplasm. The motion of the *i*-th blob was assumed to obey the overdamped Langevin equation, as follows:

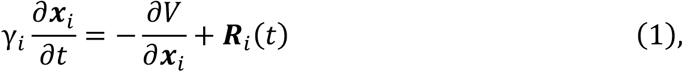

where ***x***_***i***_ = (*x*_*i*_,*y*_*i*_,*z*_*i*_) is the position of the *i*-th blob, *V* indicates the potential of the system, and *γ*_*i*_ and ***R***_*i*_ (*t*) were the coefficient of drag force and the random force working on *i*-th blob from the nucleoplasm, respectively, where *γ*_*i*_ = 6*πηr*_*i*_ with a nucleoplasm viscosity *η* and a radius of the *i*-th blob *r*_*i*_. ***R***_*i*_(*t*) was given as Gaussian white noise and satisfied ⟨***R***_*i*_ (*t*) ⟩ = 0, and ⟨*R*_*i*_ (*t*)*R*_*j*_(*s*) ⟩ = 2*γ*_*i*_*k*_*B*_*Tδ*_*ij*_*δ*(*t* − *s*), where *k*_*B*_ is the Boltzmann constant, *T* is the temperature, *δ*_*ij*_ indicates the Kronecker delta, and *δ* indicates the Dirac delta function.

The first term of the right-hand side of Eq. (1) indicates forces working on the *i*-th blob provided by the potential of the system *V*, as follows:

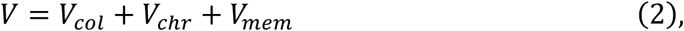

where *V*_*col*_ is the potential of the force excluding the volume effects among blobs, *V*_*chr*_ is the interaction potential of the force sustaining the shape of each chromosome, and *V*_*mem*_ is the potential of the force to confine blobs within the nuclear envelope.

The potential *V*_*col*_ was denoted by

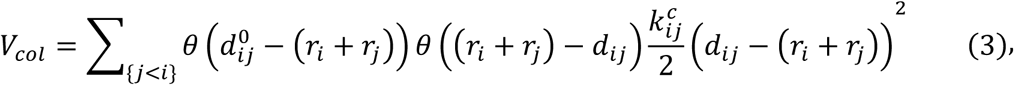

where *r*_*i*_ indicates the radius of the *i*-th blob, 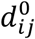 indicates the basic distance between the *i*-th and *j*-th blobs, *d*_*ij*_ = |***x***_*i*_ − ***x***_*j*_|, and *θ* is the Heaviside step function defined by the following function:

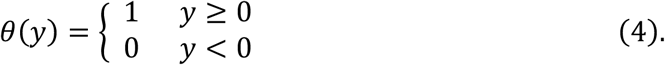

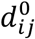 was given as the distance between the i-th and j-th blobs in the basic structure of the blob chain if the blobs belonged to the same blob chain and was given as infinity if not.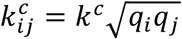 provided the coefficient of the elastic repulsion working when the *i*-th and *j*-th blobs came in contact each other, where *q*_*i*_ indicates the nondimensional parameter of rigidity of the *i*-th blob dependent on its epigenetic attribute.

The potential *V*_*bond*_ was denoted by

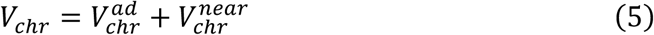

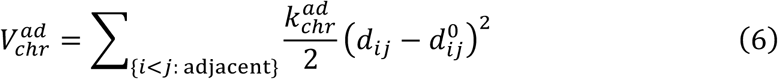

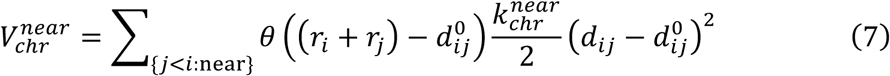

where 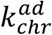 and 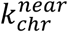 are the elastic constants sustaining the shape of the basic structure of each chromosome. The ∑ in Eq. (6) indicates the sum over the set of *i*-th and *j*-th blobs that belong to the same blob chain and adjacent to each other along the chain. The ∑ in Eq. (7) indicates the sum over the set of i-th and *j*-th blobs belonging to the same blob chain and coming in contact with each other on the basic structure of the chain but not adjacent along the chain.

The potential *V*_*mem*_ was denoted by

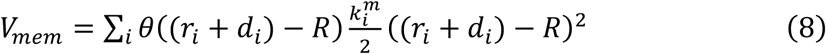

where *d*_*i*_ = |***x***_*i*_|, *R* is the radius of the nuclear envelope, and 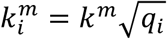 is the coefficient of the elastic repulsion working when the *i*-th blob came in contact with the nuclear envelope.

### Simulation method

To simulate the model, the time integral of the Langevin Eq. (1) was calculated numerically using the Eular-Maruyama method with a unit step of 10^−5^ s. The parameters of the nucleoplasm features were given as *η* = 0.64 *kg m*^−1^ *s*^− 1^, *k*_*B*_*T* = 4.141947 × 10^−21^ *kg m*^*2*^ *s*^− *2*^ (*T* = 300 *K*).

Because A-, B-, and M-blobs described the open flexible chromatin regions, the closed dense chromatin regions, and the region with intermediate features, respectively, M-blobs were assumed to be more rigid than A-blobs and less rigid than B-blobs. In the current study, *q*_*i*_ was assumed to be 0.1, 10.0, and 1.0 for A-, B-, and M-blobs, respectively. Other parameters were assumed to be *k*^*c*^ = 10^−4^ *kg s*^−2^, 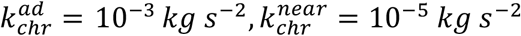, and *k*^*m*^ = 10^−4^ *kg s*^−2^. Note that physiologically precise determinations of these parameter values were difficult. Therefore, these values were chosen to be sufficiently large to avoid the crossing of two local blob chains with each other. The qualitative features of the simulation results and conclusions of the current study were not changed, even if the detail factors of these parameters were changed.

The blob chain models of homologous chromosomes were assumed to have the same basic structures, and the blob chains were randomly placed in a spherical shell with a radius of 5 μm for the initial condition of each simulation.

### Statistical analysis of simulation data

The probability distributions of the distance between two blobs containing Xic pairs were plotted with a bin interval of 1 μm, consistent with that used in previous reports of experimental results [1, 2]. Those of distances between pairs of homologous chromosomes were also plotted with the same bin interval. For both models of ES cells and 2-days cells, 30 simulations with different initial configurations of blob chains were performed. The probability distributions of these distances for each simulation were evaluated using data for blob coordinates from 9 to 36 s. For both models, the average values and 95% confidence intervals (error bars) of the probability of exhibiting each distance were calculated using 30 probability distribution profiles. The significance of the difference in the probability of exhibiting each distance between ES cell and 2-day cell models was examined by Welch’s t-test.

## Acknowledgments

Computations were partially performed on the NIG supercomputer at ROIS National Institute of Genetics.

## Author contributions

T.K. and A.A. conceived and designed the study; T.K. and A.A. conducted the mathematical model construction and simulations; T.K., M.F., and A.A. analyzed the data; T.K, M.F., and A.A. wrote the manuscript.

## References

[1] Bacher CP, Guggiari M, Brors B, Augui S, Clerc P, Avner P, et al. Transient colocalization of X-inactivation centres accompanies the initiation of X inactivation. Nat Cell Biol. 2006; 8: 293–299.

[2] Xu N, Tsai CL, Lee JT. Transient homologous chromosome pairing marks the onset of X inactivation. Science. 2006; 311: 1149–1152.

[3] Masui O, Bonnet I, Le Baccon P, Brito I, Pollex T, Murphy N, et al. Live-cell chromosome dynamics and outcome of X chromosome pairing events during ES cell differentiation. Cell. 2011; 145: 447–458.

[4] Pollex T, Heard E. Nuclear positioning and pairing of X-chromosome inactivation centers are not primary determinants during initiation of random X-inactivation. Nat Genet. 2019; 51: 285–295.

[5] Payer B, Lee JT. X chromosome dosage compensation: how mammals keep the balance. Ann Rev Genet. 2008; 42: 733–772.

[6] Nicodemi M, Prisco A. Symmetry-breaking model for X-chromosome inactivation. Phys Rev Lett. 2007; 98: 1–4.

[7] Nicodemi M, Prisco A. Self-assembly and DNA binding of the blocking factor in X chromosome inactivation. PLoS Comput Biol. 2007; 3: 2135–2142.

[8] Scialdone A, Nicodemi M. Mechanics and dynamics of X-chromosome pairing at X inactivation. PLoS Comput Biol. 2008; 4: 1–7.

[9] Scialdone A, Nicodemi M. DNA loci cross-talk through thermodynamics. J Biomed Biotechnol. 2009; 2009: 516723.

[10] Scialdone A, Cataudella I, Barbieri M, Prisco A, Nicodemi M. Conformation regulation of the X chromosome inactivation center: a model. PLoS Comput Biol. 2001; 7: e1002229.

[11] Bianco V, Scialdone A, Nicodemi M. Colocalization of multiple DNA loci: a physical mechanism. Biophys J. 2012; 103: 2223–2232.

[12] Mutzel V, Okamoto I, Dunkel I, Saitou M, Giorgetti L, Heard E, et al. A symmetric toggle switch explains the onset of random X inactivation in different mammals. Nat Struct Mol Biol. 2019; 26: 350–360.

[13] Cremer T, Cremer M, Hübner B, Silahtaroglu A, Hendzel M, Lanctôt C, et al. The interchromatin compartment participates in the structural and functional organization of the cell nucleus. BioEssays. 2020; 42: 1–18.

[14] Croft JA, Bridger JM, Boyle S, Perry P, Teague P, Bickmore WA. Differences in the localization and morphology of chromosomes in the human nucleus. J Cell Biol. 1999; 145: 1119–1131.

[15] Habermann FA, Cremer M, Walter J, Kreth G, von Hase J, Bauer K, et al. Arrangements of macro-and microchromosomes in chicken cells. Chromosome Res. 2001; 9: 569–584.

[16] Boyle S, Gilchrist S, Bridger JM, Mahy NL, Ellis JA, Bickmore WA. The spatial organization of human chromosomes within the nuclei of normal and emerin-mutant cells. Hum Mol Genet. 2001; 10: 211–219.

[17] Tanabe H, Müller S, Neusser M, von Hase J, Calcagno E, Cremer M, et al. Evolutionary conservation of chromosome territory arrangements in cell nuclei from higher primates. Proc Natl Acad Sci U S A. 2002; 99: 4424–4429.

[18] Mayer R, Brero A, von Hase J, Schroeder T, Cremer T, Dietzel S. Common themes and cell type specific variations of higher order chromatin arrangements in the mouse. BMC Cell Biol. 2005; 6: 1–22.

[19] Neusser M, Schubel V, Koch A, Cremer T, Müller S. Evolutionarily conserved, cell type and species-specific higher order chromatin arrangements in interphase nuclei of primates. Chromosoma. 2007; 116: 307–320.

[20] Solovei I, Kreysing M, Lanctôt C, Kösem S, Peichl L, Cremer T, et al. Nuclear architecture of rod photoreceptor cells adapts to vision in mammalian evolution. Cell. 2009; 137: 356–368.

[21] Koehler D, Zakhartchenko V, Froenicke L, Stone G, Stanyon R, Wolf E, et al. Changes of higher order chromatin arrangements during major genome activation in bovine preimplantation embryos. Exp Cell Res. 2009; 315: 2053–2063.

[22] Finan K, Cook PR, Marenduzzo D. Non-specific (entropic) forces as major determinants of the structure of mammalian chromosomes. Chromosome Res. 2011; 19: 53–61.

[23] Solovei I, Wang AS, Thanisch K, Schmidt CS, Krebs S, Zwerger M, et al. LBR and lamin A/C sequentially tether peripheral heterochromatin and inversely regulate differentiation. Cell. 2013; 152: 584–598.

[24] Ganai N, Sengupta S, Menon GI. Chromosome positioning from activity-based segregation. Nucleic Acids Res. 2014; 42: 4145–4159.

[25] Awazu A. Nuclear dynamical deformation induced hetero- and euchromatin positioning. Phys Rev E Stat Nonlin Soft Matter Phys. 2015; 92: 1–5.

[26] Takamiya K, Yamamoto K, Isami S, Nishimori H, Awazu A. Excluded volume effect enhances the homology pairing of model chromosomes. Nonlin Theory Appl IEICE. 2016; 7: 66–75.

[27] Seirin Lee S, Tashiro S, Awazu A, Kobayashi R. A new application of the phase-field method for understanding the mechanisms of nuclear architecture reorganization. J Math Biol. 2017; 74: 333–354.

[28] Takao K, Takamiya K, Ding DQ, Haraguchi T, Hiraoka Y, Nishimori H, et al. Torsional turning motion of chromosomes as an accelerating force to align homologous chromosomes during meiosis. J Phys Soc Jpn. 2019; 88: 1–5.

[29] Miura H, Takahashi S, Poonperm R, Tanigawa A, Takebayashi SI, Hiratani I. Single-cell DNA replication profiling identifies spatiotemporal developmental dynamics of chromosome organization. Nat Genet. 2019; 51: 1356–1368.

[30] Lieberman-Aiden E, van Berkum NL, Williams L, Imakaev M, Ragoczy T, Telling A, et al. Comprehensive mapping of long-range interactions reveals folding principles of the human genome. Science. 2009; 326: 289–293.

[31] Asakura S, Oosawa F. On interaction between two bodies immersed in a solution of macromolecules. J Chem Phys. 1954; 22: 1255–1256.

[32] Marenduzzo D, Finan K, Cook PR. The depletion attraction: an underappreciated force driving cellular organization. J Cell Biol. 2006; 175: 681–686.

[33] Zosel F, Soranno A, Buholzer KJ, Nettels D, Schuler B. Depletion interactions modulate the binding between disordered proteins in crowded environments. Proc Natl Acad Sci U S A. 2020; 117: 13480–13489.

[34] Adhikari NP, Auhl R, Straube E. Interfacial properties of flexible and semiflexible polymers. Macromol Theory Simul. 2002; 11: 315–325.

[35] Egorov SA, Milchev A, Nikoubashman A, Binder K. Phase separation and nematic order in lyotropic solutions: two types of polymers with different stiffnesses in a common solvent. J Phys Chem B. 2021; 125: 956–969.

[36] Milchev A, Egorov SA, Midya J, Binder K, Nikoubashman A. Entropic unmixing in nematic blends of semiflexible polymers. ACS Macro Lett. 2020; 9: 1779–1784.

[37] Chen CK, Blanco M, Jackson C, Aznauryan E, Ollikainen N, Surka C, et al. Xist recruits the X chromosome to the nuclear lamina to enable chromosome-wide silencing. Science. 2016; 354: 468–472.

[38] Baù D, Sanyal A, Lajoie BR, Capriotti E, Byron M, Lawrence JB, et al. The three-dimensional folding of the α-globin gene domain reveals formation of chromatin globules. Nat Struct Mol Biol. 2011; 18: 107–115.

[39] Duan Z, Andronescu M, Schutz K, McIlwain S, Kim YJ, Lee C, et al. A three-dimensional model of the yeast genome. Nature. 2010; 465: 363–367.

[40] Tanizawa H, Iwasaki O, Tanaka A, Capizzi JR, Wickramasinghe P, Lee M, et al. Mapping of long-range associations throughout the fission yeast genome reveals global genome organization linked to transcriptional regulation. Nucleic Acids Res. 2010; 38: 8164–8177.

[41] Varoquaux N, Ay F, Stafford Noble W, Vert JP. A statistical approach for inferring the 3D structure of the genome. Bioinformatics. 2014; 30: i26–i33.

