## Supporting Information for "Epigenetic-structural changes in X chromosomes promote Xic pairing during early differentiation from mouse embryonic stem cells"

**Table S1. Genomic, epigenomic, and structural features for each chromosomal region in ES cells and 2-days cells.**

See files ES_cell_model_raw.xlsx and 2-days_cell_model_raw.xlsx.

**Table S2. Genomic, epigenomic, and physical features of blobs in coarse-grained blob chain models of chromosomes from ES cells (ES cell model) and 2-days cells (2-days cell model).**

See file ES_cell_model_cg.xlsx and 2days_cell_model_cg.xlsx.

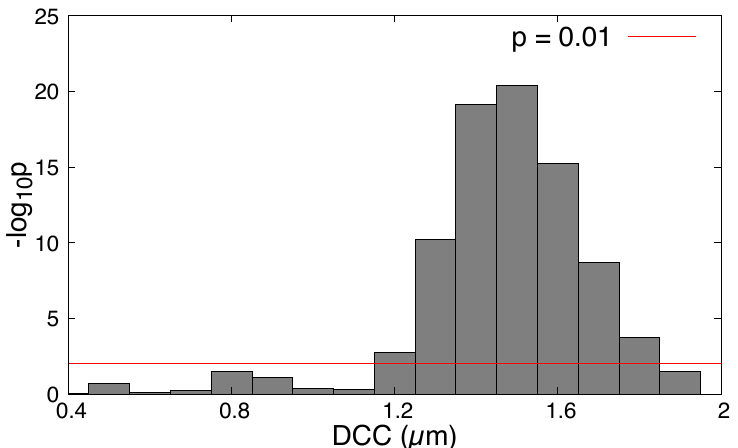

**Fig S1. Distributions of *p* values (-log_10_ *p*) for tests between RPD values.** Distributions of *p* values (-log_10_ *p*) from Welch’s t-tests between RPD values obtained for two neighboring DCC values.

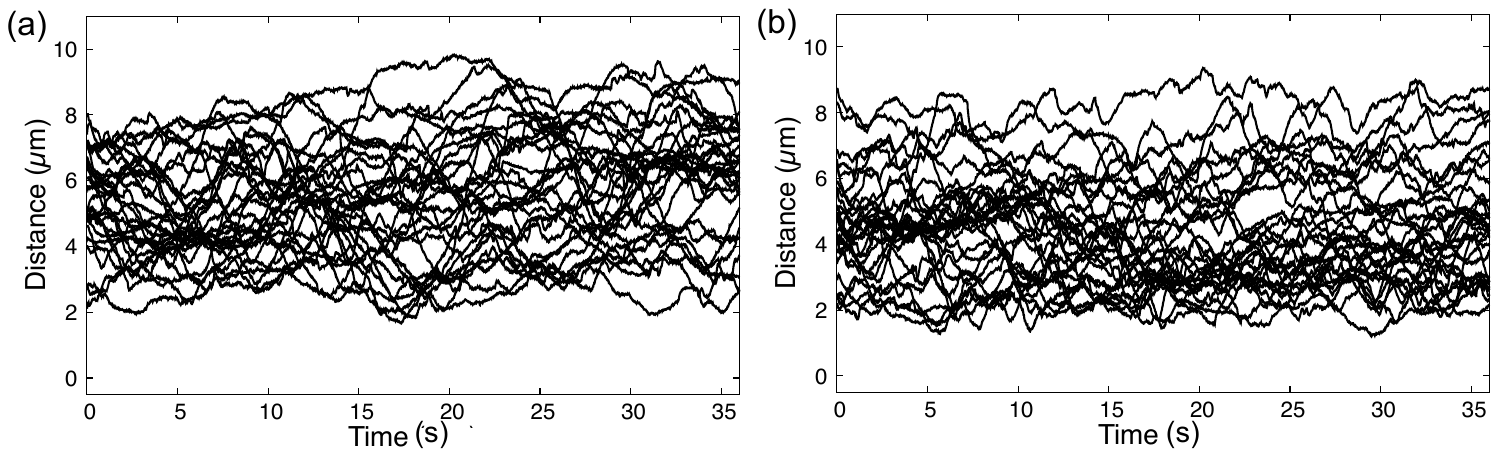

**Fig S2. Time course of distances between Xic pairs.** Simulation results of temporal changes in distances between two Xic pairs obtained from 30 different initial conditions in the ES cell model (a) and 2-days cell model (b).

**Fig S3. Probability distributions of distances between homologous chromosomes.** Averages and 95% confidence intervals (error bars) of probability distributions of distances between pairs of homologous chromosomes in the ES cell and 2-days cell models. Averages and 95% confidence intervals were evaluated using the results from 30 simulations for each model. Distances with asterisks indicate that the differences in probability values between the two models were significant (*p* < 0.05 by Welch’s t-test; see also Fig S3).

**Fig S4. Distributions of *p* values (-log_10_ *p*) for tests of probability distributions for distances between homologous chromosomes.** Distributions of *p* values (-log_10_ *p*) from Welch’s t-tests between probability values obtained in the ES cell and 2-days cell models at each distance for pairs of homologous chromosomes (Fig S2).

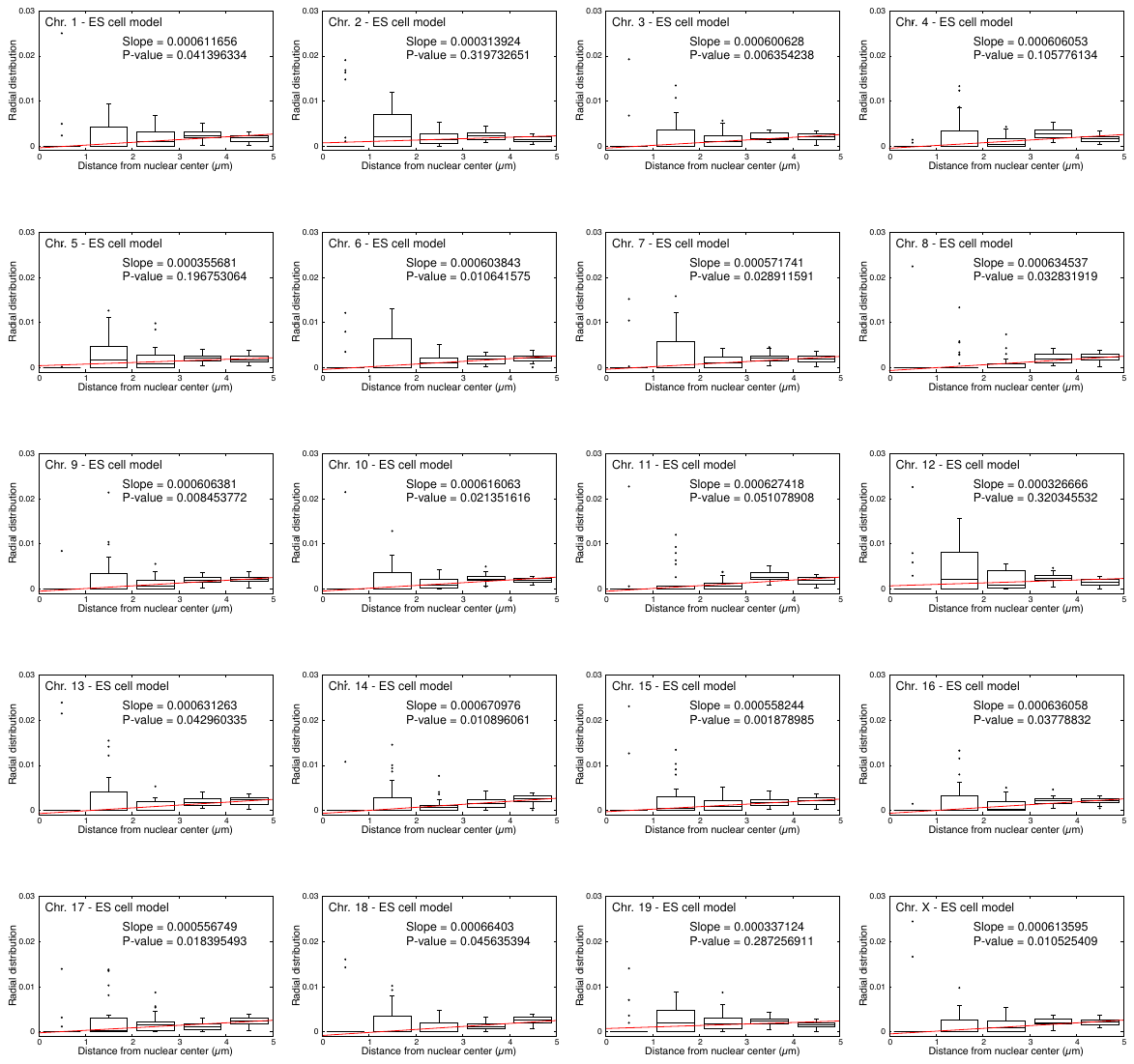

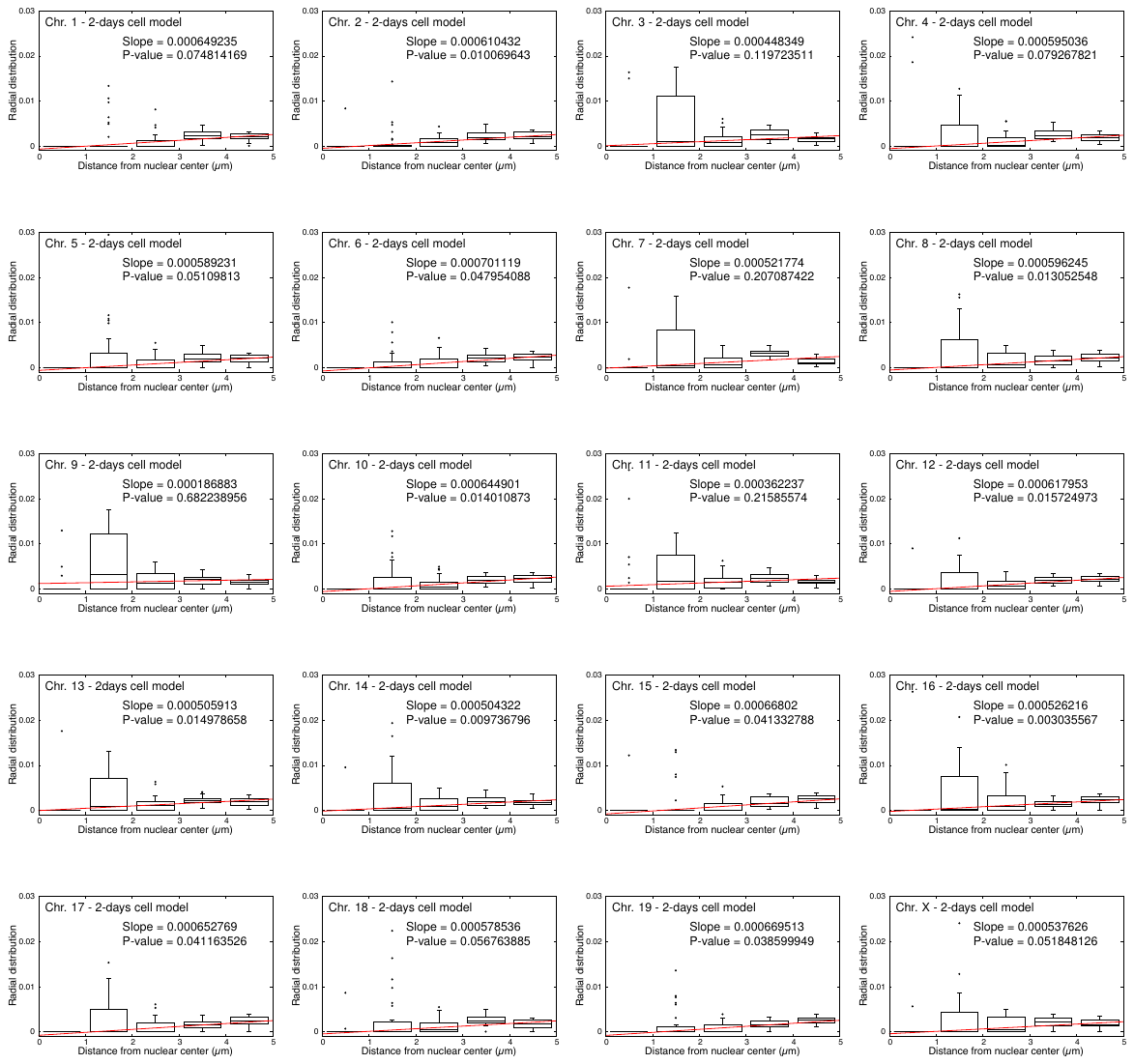

**Fig S5. Radial distribution functions of Xic pairs or chromosomes.** Box plots of values of radial distribution functions from the nuclear center to blobs containing Xic pairs and chromosomes in ES cell and 2-days cell models. Radial distribution functions were defined as (the frequency distribution of distances from the center of nucleus to Xic or each chromosome) / ($4\pi$ [distance]^2^). The box plots of radial distributions for both ES and 2-days cell models were plotted using data from 30 simulations of these models. The distance-dependent features of these radial distributions were evaluated by slopes and *p* values evaluated by linear regression analysis between the distance and the median value of radial distribution for each distance.

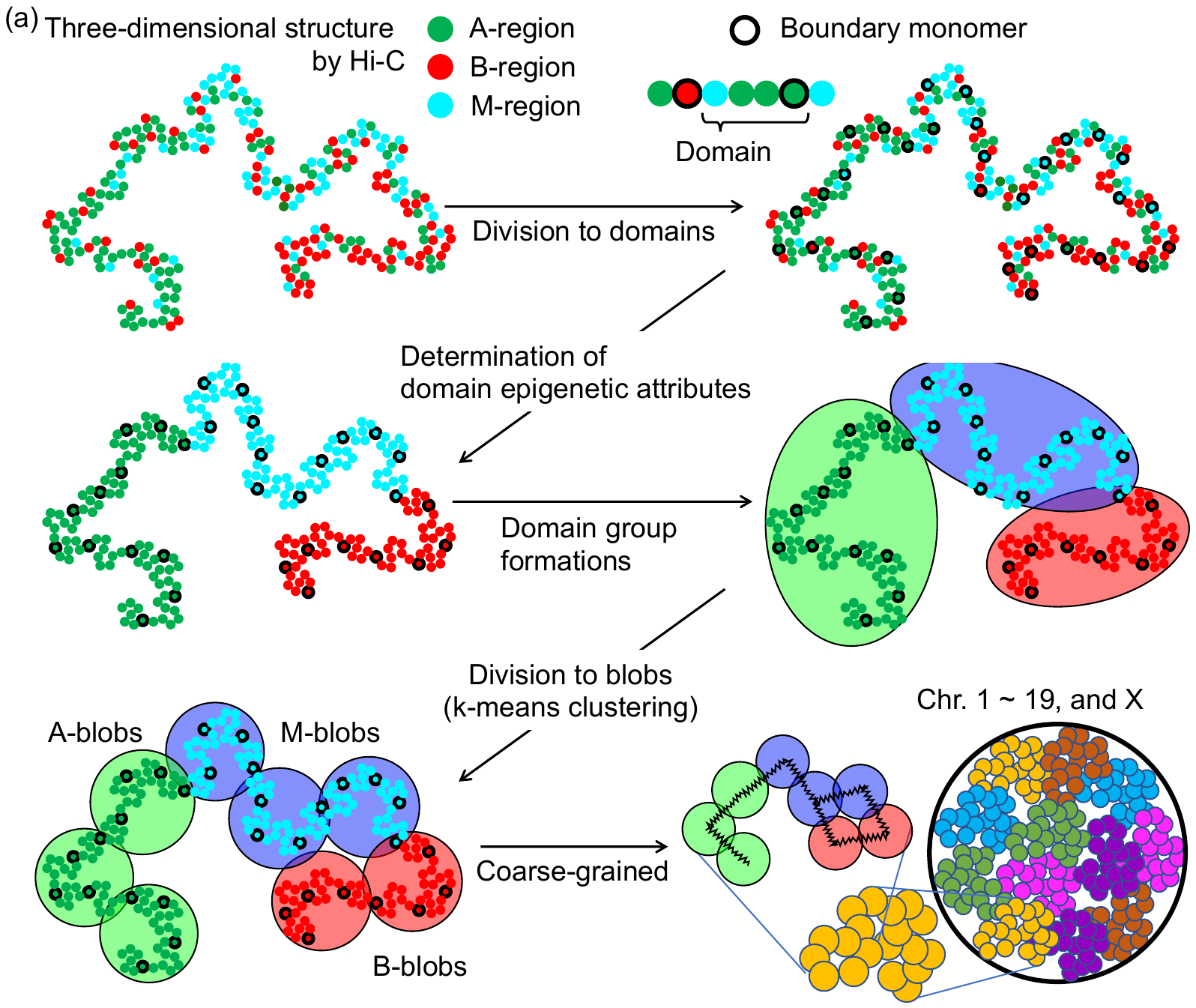

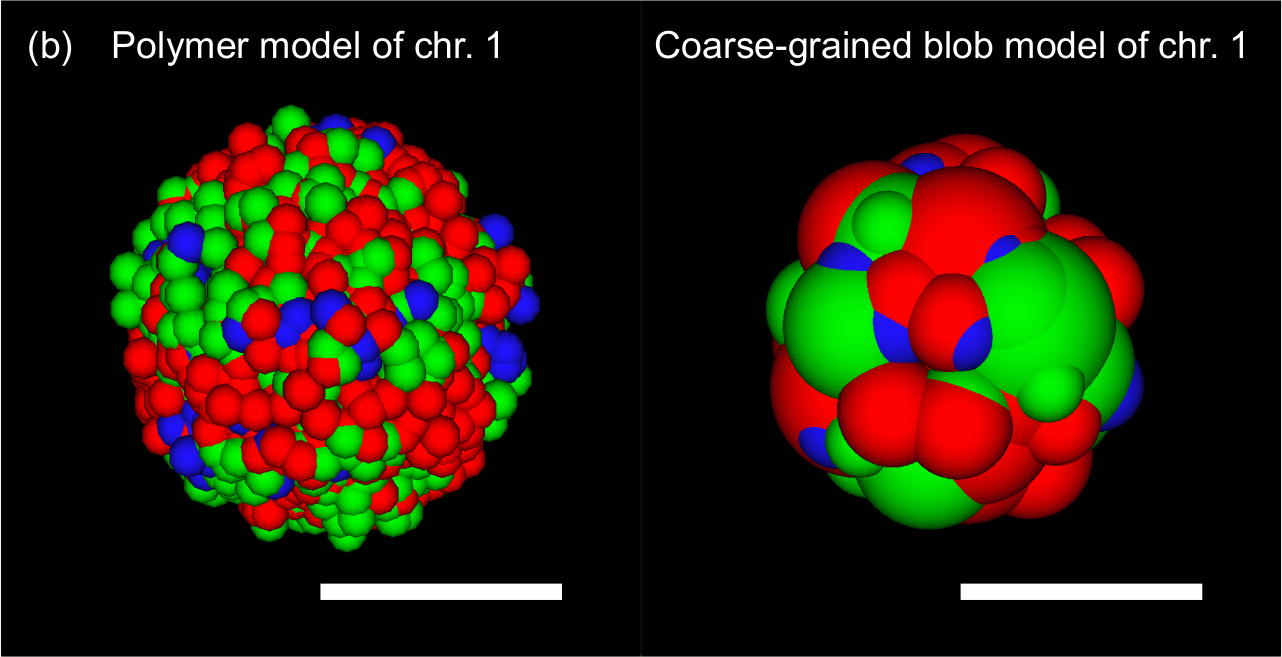

**Fig S6. Illustration of the coarse-grained model construction scheme.** (a) This workflow was illustrated using virtual sample data to show each process more clearly. In this example, CDN_n_ was assumed as four regions for k-means clustering. Each pair of overlaid blobs was connected by an elastic spring with a natural length equal to the distance between their centers (right bottom). Detailed explanations of each modeling step are given in the Methods section. (b) Illustrations of the polymer and blob models, using chromosome 1 in ES cells as an example. Scale bar: 2 μm.

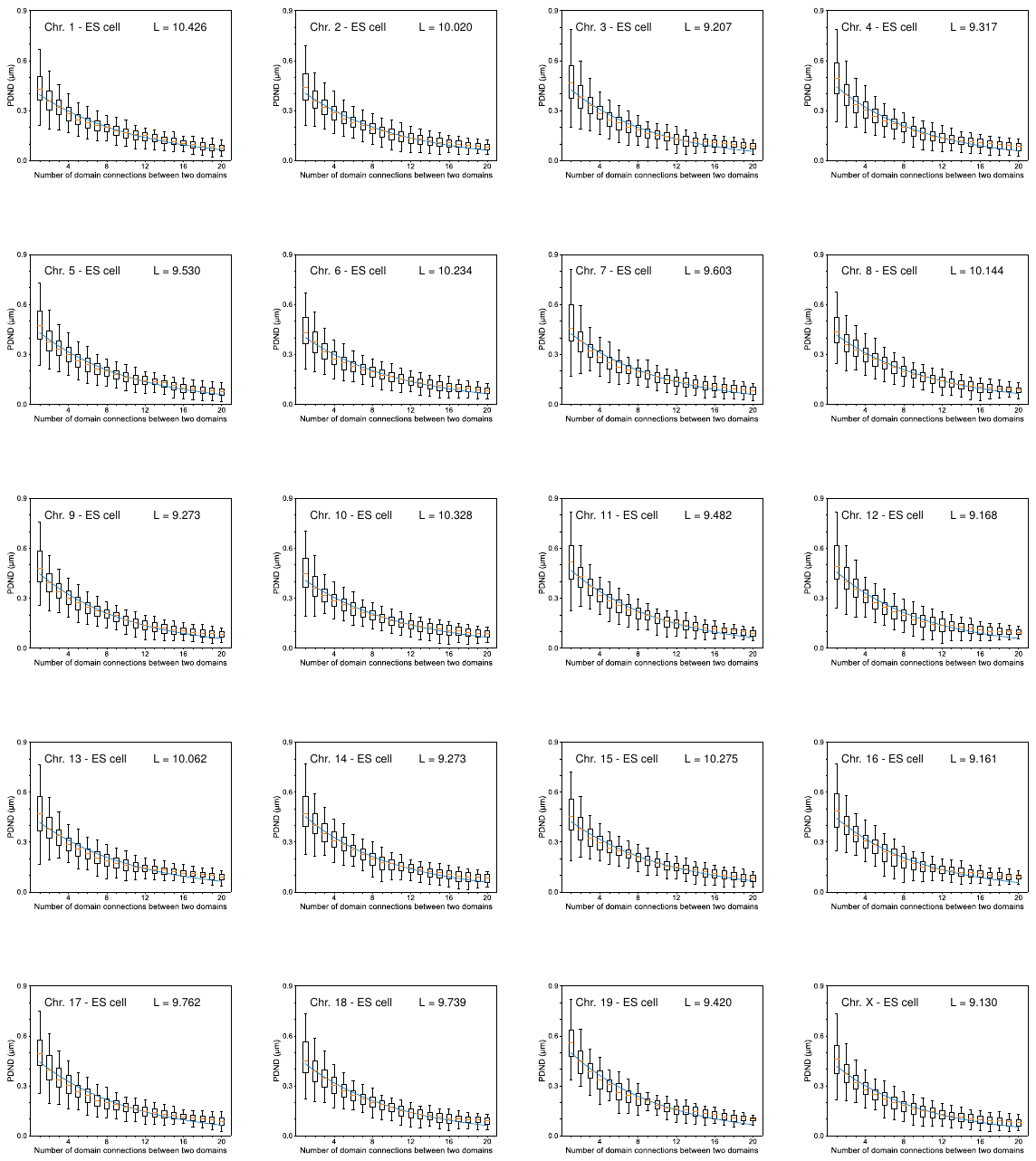

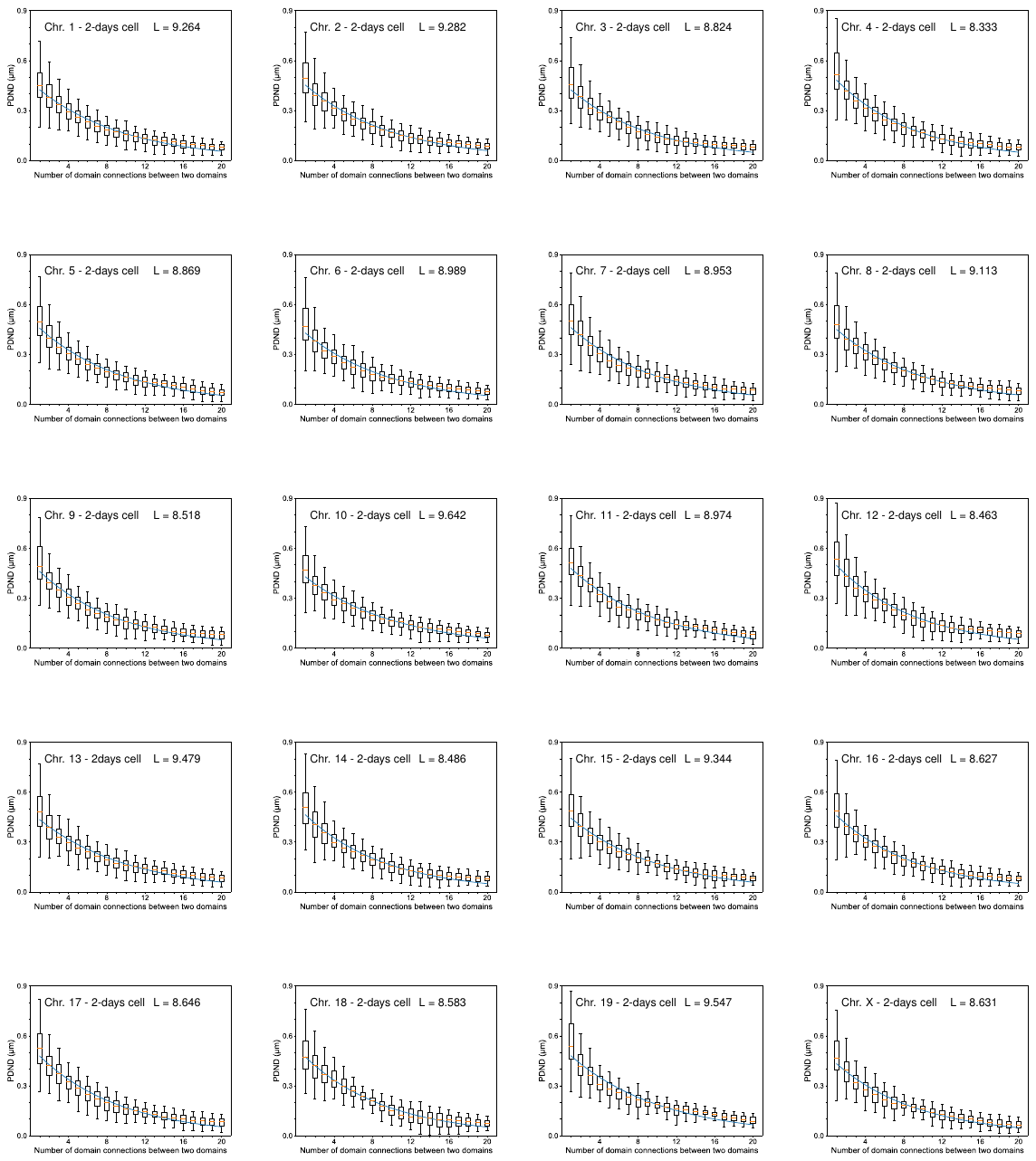

**Fig S7. PDND to determine the characteristic number of domains (CND_n_).** Box plots of PDND as a function of the number of domain connections between two domains for all chromosomes in ES cells and 2-days cells obtained using the polymer model. Each average of PDND was estimated by PDND values within 95% confidence intervals. Curves (blue) are fitted averages of PDND that were assumed as exp(-[number of domain connections between two domains] / L), where L obtained from n-th chromosome gives CND_n_.
